# The effect of perturbation magnitude on lower limb muscle activity during reactive stepping using functional data analysis

**DOI:** 10.1101/2020.12.17.423261

**Authors:** Tyler M. Saumur, Jacqueline Nestico, George Mochizuki, Stephen D. Perry, Avril Mansfield, Sunita Mathur

## Abstract

This study aimed to determine the effect of perturbation magnitude on stance and stepping limb muscle activation during reactive stepping using functional data analysis. Nineteen healthy, young adults responded to 6 small and 6 large perturbations using an anterior lean-and-release system, evoking a single reactive step. Muscle activity from surface electromyography was compared between the two conditions for medial gastrocnemius, biceps femoris, tibialis anterior, and vastus lateralis of the stance and stepping limb using functional data analysis. Stance limb medial gastrocnemius and biceps femoris activation increased in the large compared to small perturbation condition immediately prior to foot-off and at foot contact. In the stepping limb, significant increases in medial gastrocnemius, biceps femoris, and tibialis anterior activity occurred immediately prior to foot-off during the large perturbations. Similar to the stance limb, medial gastrocnemius and biceps femoris activity significantly increased during and following foot contact in the large, compared to small, perturbation condition. Lastly, vastus lateralis activity significantly increased for large, compared to small, perturbations during foot-off and immediately following foot contact. These findings highlight lower limb muscle activity modulation associated with perturbation magnitude throughout reactive stepping and the additional benefit of implementing functional data analysis to study reactive balance control.

## 1. Introduction

Reactive balance control requires the use of different sensorimotor strategies to return the centre of mass within the base of support after a balance perturbation. Specifically, the timing and magnitude of muscle activity following a perturbation are commonly investigated in the context of reactive balance control (Cronin et al., 2013; Dietz et al., 1989; Eng et al., 1994; Forghani et al., 2017; Freyler et al., 2015; Pijnappels et al., 2005a, 2005b; Thelen et al., 2000). Many features can affect the neuromuscular response following a perturbation including body position, the surrounding support environment, and the speed, magnitude and predictability of the perturbation (Carr and Shepherd, 2003). During feet-in-place responses in unilateral and bilateral stance, amplitude of muscle activation is scaled to perturbation size (Forghani et al., 2017; Freyler et al., 2015; Mochizuki et al., 2004). While feet-in-place strategies are effective for smaller perturbations, they are not appropriate for large balance perturbations that can lead to a fall. Change-in-support strategies, such as reactive steps, are more prevalent in novel scenarios and are the only strategies that can be used to successfully regain balance after large-magnitude perturbations (Maki and McIlroy, 1997).

Forward reactive stepping, which is the most common direction of a loss of balance (Stevens et al., 2014), requires the sequential activation of the stance and stepping leg muscles to regain stability. During a trip, the posterior muscles of the stance limb are typically activated first, followed by the anterior musculature (Pijnappels et al., 2005b). The hamstrings and plantar flexors of the stance limb are important for decelerating forward momentum following the perturbation (Pijnappels et al., 2005a). Similarly, in the stepping limb, the plantar flexors and hamstrings are activated early to facilitate hip and ankle extension (Pijnappels et al., 2005a). During a trip, activating the hamstrings prior to quadriceps facilitates knee flexion before hip flexion to carry the limb over an obstacle if present (Eng et al., 1994). Conversely, dorsiflexor activation is variable, particularly in the stance limb, during trip-evoked reactive stepping (Eng et al., 1994; Pijnappels et al., 2005b). Similar findings have also been observed in these muscle groups during early reactive stepping responses to platform perturbations (prior to foot-off; Chvatal et al., 2011; Wang et al., 2016). In terms of motor unit recruitment, people who can respond to a perturbation with a single step have been shown to activate a larger proportion of their motor unit pool for the stepping leg compared to those to take multiple steps (Cronin et al., 2013), highlighting the need for greater muscle activity to successfully regain stability in a single step.

One method to understand the importance of modulating muscle activity is to modify perturbation magnitude. To date, there is limited work exploring the effect of perturbation magnitude on muscle activity during reactive stepping. Thelen et al. (2000) investigated changes in mean muscle activity with different perturbation magnitudes and found that perturbation magnitude increased activity of gastrocnemius, rectus femoris, and biceps femoris of the stance limb and tibialis anterior, vastus lateralis, and rectus femoris of the stepping limb. However, this study did not focus on differences in muscle activity throughout the stepping response, limiting our understanding of the functional specificity to the muscle activity changes. Understanding how muscle activity is modified in response to different perturbation magnitudes throughout the different reactive stepping phases can further our understanding of how reactive stepping can effectively maintain balance in different contexts.

Reactive stepping involves multiple phases. Typical analyses used to study muscle activity, such as multivariate analyses, may separate electromyographic (EMG) data based on the response phases of reactive stepping (e.g. swing phase, landing phase). These techniques often analyze EMG onsets or magnitudes in a discrete manner by comparing peak muscle activity or average muscle activity over a pre-defined time period (Cronin et al., 2013; Pijnappels et al., 2005b; Thelen et al., 2000). However, separating the stepping response into different phases that are analyzed in isolation limits our ability to understand the variability and smoothness of EMG time series data continuously and can result in missing important differences. Functional data analysis (FDA) is a statistical approach that continuously assesses differences in data by transforming signals into curves comprised of small basis functions (Park et al., 2017; Ramsay and Silverman, 2005). The weights of these functions can then be compared statistically using a functional t-test or functional analysis of variance. Thus, FDA has the advantage of being able to evaluate the smooth functional behaviour of data to further model and understand the signal throughout an entire response in an easily interpretable manner. FDA has been previously applied to study differences in muscle activity during forward lunging (Hopkins et al., 2013) and it may have utility for detecting condition differences in reactive stepping muscle activity.

Accordingly, this study aimed to determine the effect of perturbation magnitude on muscle activation of the stance and stepping limbs during reactive stepping using FDA. We hypothesized that the larger perturbation magnitude, compared to the smaller magnitude, would result in increased muscle activation of the most relevant muscles for each leg (i.e. plantar flexors and knee flexors of the stepping leg during foot-off and foot contact; knee extensors during the swing phase of stepping leg; and plantar and knee flexors of the stance leg during foot-off).

## 2. Methods

### 2.1. Participants

Participants in this study have been described in detail previously (Saumur et al., 2020) where they participated in a test-retest reliability study. EMG was recorded during the first session and used in the present study. Young, healthy participants from the Greater Toronto Area were recruited to participate in this study. Participants were included if they met the following criteria: between the ages of 20 and 35 years, no self-reported walking difficulties, able to understand English, no neurological or musculoskeletal disorders, normal or corrected to normal vision, and no conditions that limit the ability to complete daily activities. Participants provided written informed consent and research ethics board approval was obtained from Sunnybrook Health Sciences Centre (#017-2015) and the University of Toronto (#31805).

### 2.2. Task procedures

Participants underwent reactive balance testing using a lean-and-release system (Figure 1; Inness et al., 2015). Initially, participants were given a demonstration of the lean-and-release perturbation to ensure they felt comfortable with the task. Participants stood on a platform with embedded force plates and were fitted with a harness to ensure safety. Participants were connected to a second tether at the rear wall, released at a random time point by the investigator, and instructed to “recover balance in as few steps as possible” with the specified leg. An initial small test perturbation was performed by the participant to remove first trial effects and familiarize them with the type of balance loss. Participants then performed two blocks (one for each leg) of six perturbation trials randomized to perturbation magnitude, resulting in three trials at each perturbation magnitude for each leg. The block order was randomized between participants. Small (~8-10% of body weight) and large (~13-15% of body weight) magnitude perturbations were implemented in this study and magnitude size was increased by lengthening the tether (lean angle) accordingly. EMG and force plate data were monitored online to ensure that no anticipatory offloading occurred prior to the perturbation.

**Figure 1.**
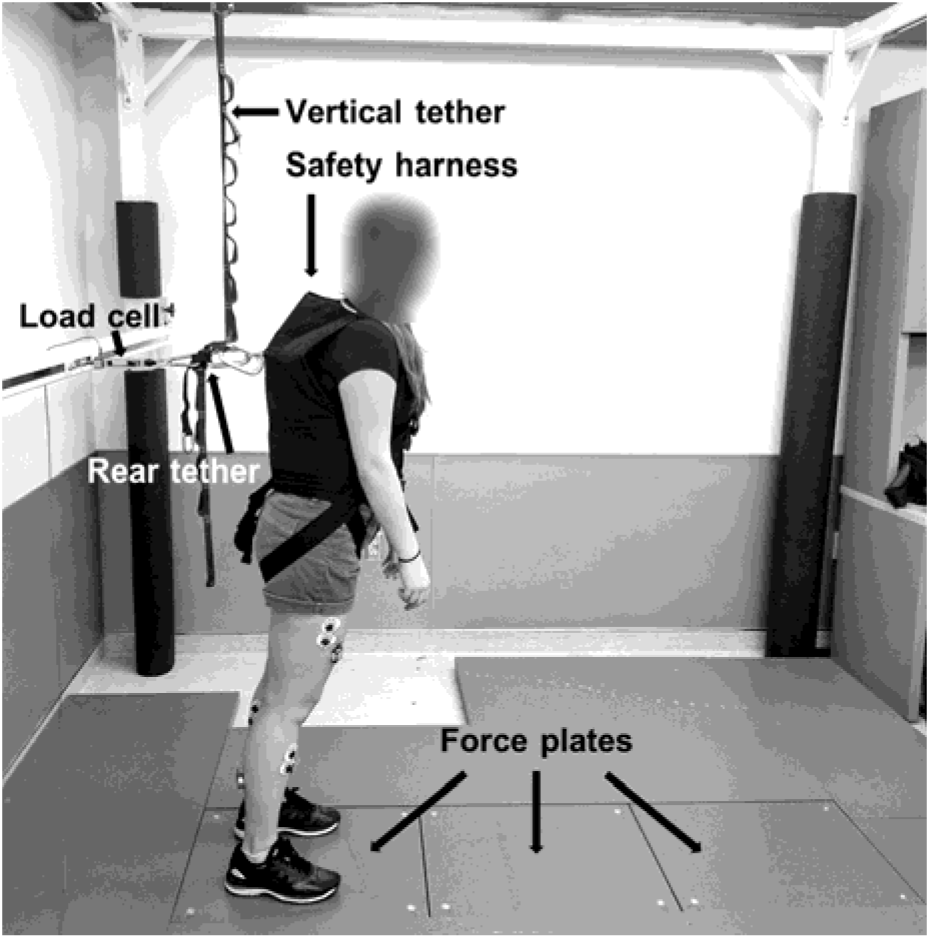
Example of experimental setup.

### 2.3 Electromyography (EMG)

Surface EMG was collected bilaterally from the following muscles: medial gastrocnemius (MG), biceps femoris (BF), tibialis anterior (TA), and vastus lateralis (VL) using 1.375” self-adhesive circular Ag-AgCl recording electrodes (Kendall 530 Series, Cardinal Health Inc., Dublin, Ohio, USA). A bipolar electrode configuration was used with an interelectrode distance of 20 mm; electrodes were oriented in line with the direction of the muscle fibres. Skin was prepared and cleaned with abrasive preparation gel (NuPrep, Weaver and Company, Aurora, Colorado, USA) and an alcohol swab prior to electrode placement. EMG signals were sampled wirelessly at 1500 Hz using the TeleMyo DTS system (Noraxon, Scottsdale, Arizona, USA) and transmitted to a computer using the G2 Analog Output receiver (Noraxon, Scottsdale, Arizona, USA). All EMG data were normalized to maximal voluntary isometric contractions (MVICs) obtained for each muscle using the HUMAC NORM dynamometer on the preferred stepping limb (CSMi Medical Solutions, Stoughton, Massachusetts, USA) as limb dominance has little effect in young, healthy adults (McCurdy and Langford, 2005). Participants performed 5 second MVICs for knee extension and ankle plantarflexion and dorsiflexion in the supine position and knee flexion in the prone position to not introduce EMG noise. All joint angles for testing were determined based on the muscle lengths that maximize muscle activity and torque generation (Arampatzis et al., 2006; Dalton et al., 2013; Kulig et al., 1984; Onishi et al., 2002; Sale et al., 1982).

Offline filtering was performed using Spike2 software (Spike2 Version 7.17, Cambridge Electronic Design, Cambridge, UK). EMG data for each trial were full wave rectified and low pass filtered at 80 Hz to create a linear envelope. The data were then scaled based on MVIC EMG activity. The highest mean EMG value obtained over a 500 msec period from all MVIC trials was used for normalizing each muscle group. A root mean square algorithm with a 50 msec window was then applied for each trial (Hopkins et al., 2013, 2012). Data were time normalized such that perturbation onset occurred at 0% and foot contact occurred at 33% of the trial duration for all participants. Stance and stepping limb EMG data were analyzed for all four muscles. Perturbation onset was defined as < 5 N of force on the load cell, and foot contact occurred when the stepping force plate ground reaction forces exceeded 5 N (Saumur et al., 2020). Foot-off time was determined as the time when the mediolateral centre of pressure was 20% of the maximum slope when shifting towards the stepping limb (Kurz et al., 2013; Saumur et al., 2020). Trials were excluded if participants took more than one step to respond to the perturbation. EMG responses during balance testing were excluded at the time of analysis if there was anticipatory activation during or prior to perturbation onset (activation exceeding 10% of the peak EMG response for the trial).

### 2.4. Statistical analysis

All data are reported as means (standard deviations) unless otherwise stated. FDA was conducted in RStudio (Version 1.3.959). Warping functions were fitted to the data so that all trials had the same start and end points. Once the data were time registered, it was smoothed using B-splines with 19 basis functions (Park et al., 2017). To highlight the differences between these functions, the difference between the pairwise comparison functions (large perturbation condition – small perturbation condition) were graphed along with the 95% confidence intervals (Figure 2). Differences between conditions were deemed statistically significant when the 95% confidence interval did not cross zero (Park et al., 2017). Differences between large and small perturbations are reported based on the lower bounds of the 95% confidence intervals.

**Figure 2.**
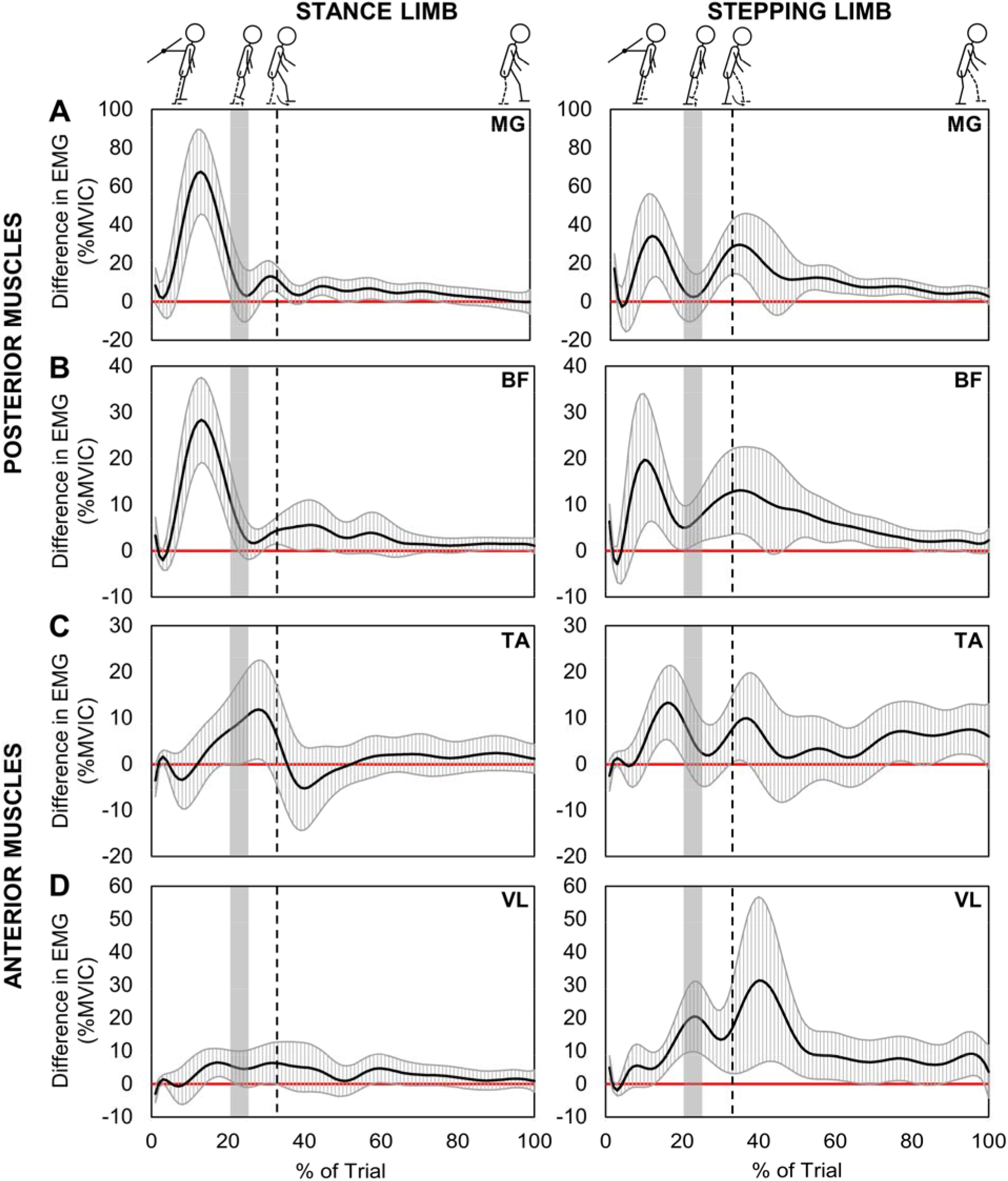
Results of functional data analyses for all 8 muscles (solid black line-mean, grey hatched band-95% CI). These graphs represent the differences of large perturbation condition minus small perturbation condition. Significant differences occur where the 95% confidence intervals (CI) do not touch the red zero line. The grey bar represents the approximate time window of foot-off (calculated from the mean foot-off time and trial time for all participants and conditions plus or minus 2 standard deviations) and the dotted vertical line represents foot contact. Medial gastrocnemius (MG), biceps femoris (BF), tibialis anterior (TA), and vastus lateralis (VL).

## 3. Results

Nineteen participants took part in this study (27.6 (3.0) years; 9 men and 10 women) (Saumur et al., 2020). A summary of the number of trials used for analysis for each muscle can be found in Table 1. Approximately 93% of the data were retained; reasons for trial removal during data analysis included: muscle pre-activation (2.7%), movement following the reactive step that was not a result of the perturbation or the inability to maintain stability on the force plates following the perturbation (2.6%), and EMG issues (i.e. EMG unit stopped recording) or noise (1.6%).

**Table 1.** Total number of trials included in analysis for each muscle

| Muscle | Total trials included (%) |
| --- | --- |
| **Stance Limb** |  |
| Biceps femoris | 211 (93%) |
| Medial gastrocnemius | 217 (95%) |
| Tibialis anterior | 217 (95%) |
| Vastus lateralis | 205 (90%) |
| **Stepping Limb** |  |
| Biceps femoris | 207 (91%) |
| Medial gastrocnemius | 214 (94%) |
| Tibialis anterior | 218 (96%) |
| Vastus lateralis | 208 (91%) |

### 3.1. Muscle activation of the stance limb

The FDA revealed several differences in stance limb muscle activation between the large and small perturbation conditions (Figure 2). All significant differences resulted in greater muscle activation during the large perturbation condition. Most notably, these changes were observed in the posterior muscles (MG and BF) prior to foot-off (Figures 2A and B). The largest significant difference observed among all muscles for both limbs occurred in the stance limb MG prior to foot-off, with muscle activity increasing by 45% of MVIC in the large perturbation condition. There were also smaller significant differences (<5% of MVIC) in stance limb MG activity during foot contact and periodically throughout stabilization following foot contact.

Similar to MG, the largest difference in stance limb BF muscle activity occurred prior to foot-off (increase of 19% of MVIC). A small significant increase in BF muscle activity also occurred around the time of foot contact. For stance limb TA and VL, there were small significant increases (<3% MVIC) in muscle activation during the swing phase and prior to foot-off, respectively (Figures 2C and D).

### 3.2. Muscle activation of the stepping limb

Several significant differences were found between conditions for the stepping limb with all significant differences resulting in greater muscle activation during the large perturbation condition. For MG, BF, and TA (Figures 2A, B and C), muscle activity was significantly higher prior to foot-off (increase of 5-13% of MVIC) in the stepping limb. Conversely, VL activity increased by 10% of MVIC during the swing phase and 7% of MVIC immediately following foot contact (Figure 2D). MG and BF also showed a significant increase in muscle activity around the time of foot contact and this was maintained into the later portion of stabilization. TA had slightly higher activation during foot contact and during the last 10% of the trial.

## 4. Discussion

This study explored the effect of perturbation magnitude on lower limb muscle activity using FDA to continuously assess differences in the EMG signals. FDA revealed that all differences resulted in greater muscle activation in the large perturbation condition compared to the small perturbation condition. For the stance limb, the following significant differences were observed: (1) increased MG and BF activity immediately prior to foot-off and to a lesser extent at foot contact; and (2) a small increase in TA and VL activity during the swing phase and prior to foot-off, respectively. In the stepping limb, analysis revealed a: (1) significant increase in MG, BF, and TA activity immediately prior to foot-off; (2) significant increase in MG and BF activity during and following foot contact; and (3) significant increase in VL activity at foot-off and immediately following foot contact. These findings suggest specific perturbation-induced modulations in muscle activity of both the stance and stepping limb during reactive stepping.

### 4.1. The modulation of posterior muscle activity for deceleration, propulsion & braking forces

The appropriate scaling of muscle activity during a perturbation is important to regain stability following a loss of balance (Cronin et al., 2013). In this study, we found magnitude-specific modulation in the muscles of both the stance and stepping limb throughout reactive stepping. While we had hypothesized that differences in posterior musculature would occur at foot-off, in general, the large perturbation magnitude resulted in increased activation of the posterior musculature immediately prior to foot-off for both the stance and stepping limb. During tripping, the MG and BF of the stance limb are important in the early phases of the response for generating large ankle plantarflexion and large hip extension moments, respectively (Pijnappels et al., 2005a). The combination of these moments is an attempt to reduce forward angular momentum to promote a successful recovery (Pijnappels et al., 2005a, 2004). After foot-off, the hip abductors and plantarflexors have been shown to accelerate spine flexion and hip extension of the stepping limb (Graham et al., 2014). However, our results suggest that there is little modulation of MG and BF during the swing phase as perturbation magnitude increases.

In the stepping limb, the rapid, early activation of MG and BF is responsible for centre of mass propulsion through ankle flexion and hip extension (Graham et al., 2017). We observed increased muscle activity in the stepping limb posterior muscles which may be required for a faster reactive step during the larger perturbation condition. Indeed, faster swing time and shorter step lengths have been observed with increasing perturbation magnitude (Thelen et al., 2000). Interestingly, when comparing the differences which were observed between limbs, the lower bounds of the difference in MG and BF muscle activity between the large and small perturbation condition were substantially larger in the stance limb than the stepping limb (MG: 45% compared to 13% of MVIC, BF: 19% compared to 6% of MVIC); this may suggest that the central nervous system prioritizes the reduction of forward momentum, rather than increasing force generation to propel the stepping limb when perturbation magnitude increases.

During the later phases of the reactive step, all peak differences observed in MG and BF for both the stance and stepping limb occurred at foot contact aligning with our hypotheses and highlighting the increased importance of modulating muscle activity to absorb and brake the body’s forward momentum produced during the large perturbation condition upon contact (Thelen et al., 2000). These findings contrast those of Thelen et al (2000), in which there was only a significant increase in mean dorsiflexor and knee extensor muscle activity with increased perturbation magnitude. This highlights the advantage of using FDA to analyse the EMG time series and reveal effects of perturbation magnitude on muscle activity. Stepping limb MG activity demonstrated larger differences during foot contact compared to the stance limb (15% compared to 5% of MVIC); this may indicate an increased need to prioritize stiffening the stepping limb ankle joint compared to absorbing impact with the stance leg. While the posterior muscles of the leg act about different joints, their increased activation during large perturbations appears to be important for decelerating body momentum, propelling the stepping limb forward, and applying braking forces at foot contact.

### 4.2. The modulation of anterior muscle activity for foot-off time and stabilization

Aligning with our hypothesis and the limited role of the TA in tripping (Eng et al., 1994; Pijnappels et al., 2005b), our results showed little modulation in TA activity with increased perturbation magnitude. A small significant increase in TA activity during the large perturbation condition was observed in the stance limb during the swing phase. In contrast to our hypothesis, a significant modulation of swing limb TA activity was observed immediately prior to foot-off. This may be due to differences in perturbation methods. In the previous studies, obstacles were used to cause a trip during the swing phase of the gait cycle (Eng et al., 1994; Pijnappels et al., 2005b). As participants had already elevated their foot off the ground, TA may not have a large role in the reaction. Conversely, a significant increase in mean TA activity from small to large lean-and-release perturbations has been observed previously (Thelen et al., 2000); this modulation likely facilitated rapid foot-off by increasing muscle activation to dorsiflex the foot of the stepping limb and aligns with the timing of activation in the present study.

In VL, the peak differences between perturbation magnitudes were observed at foot-off and following foot contact in the stepping limb. The significant increase in VL activity at foot-off aligns with its role in pushing off during reactive and volitional stepping (Chvatal et al., 2011; Wang et al., 2016; Winter and Yack, 1987). The peak difference in muscle activity immediately following foot contact aligns with previous literature that highlights the role of VL in braking during perturbed walking, running, and stance (Chan et al., 2020; Ellis et al., 2014; Graham et al., 2017). Therefore, it appears that increased TA and VL activity prior to, and during foot-off facilitated faster and stronger push offs from the stepping limb. In addition, VL appears to be an important muscle for stabilizing the centre of mass following foot contact.

### 4.3. Limitations

First, step timing was estimated using force plates rather than kinematics. Particularly, foot-off was estimated from a single force plate using the slope of the centre of pressure (Kurz et al., 2013). However, the step times in this study are similar to previous studies using kinetic and kinematic methods to determine step timing (Lakhani et al., 2011; Melzer et al., 2007; Singer et al., 2019; Thelen et al., 2000). A second limitation to our study is that MVICs were performed at a pre-specified angle based on optimal muscle tissue dynamics. These angles do not necessarily reflect the muscles’ functional angles during reactive stepping and therefore may not represent motor unit recruitment observed during dynamic activities.

### 4.4. Conclusions

This is the first study to our knowledge to use FDA to explore muscle activation changes resulting from increasing perturbation magnitude during reactive stepping. We found that the modulation of posterior muscle activity was likely important for increased deceleration (stance limb) and propulsion and braking forces (stepping limb) during the large perturbation magnitude condition, based on the timing of activation changes. Conversely, increased activation of the anterior leg muscles appears to have a functional role – particularly in the stepping limb – in foot-off and stabilization following foot contact. FDA may have utility for various applications in studying reactive balance control to compare differences between subject groups or conditions and warrants further investigation.

## Acknowledgements

The authors would like to thank Dr. J. Ty Hopkins and Seunguk Han of Brigham Young University for sharing their functional data analysis code. The investigators would also like to thank all participants for their time. TMS was funded by the Ontario Graduate Scholarship, Toronto Rehabilitation Institute Student Scholarship, and Peterborough KM Hunter Charitable Foundation Graduate Award.

